# Subunit epsilon of *E. coli* F_1_F_o_ ATP synthase attenuates enzyme activity by modulating central stalk flexibility

**DOI:** 10.1101/2020.09.30.320408

**Authors:** Meghna Sobti, James L. Walshe, Yi C. Zeng, Robert Ishmukhametov, Alastair G. Stewart

## Abstract

F_1_F_o_ ATP synthase functions as a biological rotary generator that makes a major contribution to cellular energy production. Proton flow through the F_o_ motor generates rotation of the central stalk, inducing conformational changes in the F_1_ motor that catalyzes ATP production via flexible coupling. Here we present a range of cryo-EM structures of *E. coli* ATP synthase in different rotational and inhibited states observed following a 45 second incubation with 10 mM MgATP. The structures generated describe multiple changes that occur following addition of MgATP, with the inhibitory C-terminal domain of subunit ε (εCTD) disassociating from the central stalk to adopt a condensed “down” conformation. The transition to the εCTD down state increases the torsional flexibility of the central stalk allowing its foot to rotate by ∼50°, with further flexing in the peripheral stalk enabling the *c*-ring to rotate by two sub-steps in the F_o_ motor. Truncation mutants lacking the second helix of the εCTD suggest that central stalk rotational flexibility is important for F_1_F_o_ ATP synthase function. Overall this study identifies the potential role played by torsional flexing within the rotor and how this could be influenced by the ε subunit.

## Main Text

A key component in the generation of cellular metabolic energy is the F_1_F_o_ ATP synthase, a biological rotary motor that converts proton motive force (pmf) to adenosine tri- phosphate (ATP) in both oxidative phosphorylation and photophosphorylation^1-3^. The enzyme is comprised of two rotary motors, termed F_1_ and F_o_, that are coupled together by two stalks: a central “rotor” stalk and a peripheral “stator” stalk. The F_o_ motor spans the membrane and converts the potential energy from the pmf into mechanical rotation of the central rotor that, in turn, drives conformational changes in the catalytic F_1_ motor subunits that generate ATP from ADP and inorganic phosphate (P_i_)^4,5^. *E. coli* contains the simplest form of F_1_F_o_ ATP synthase, with only eight different types of subunit, and has therefore been used extensively as a model system for ATP synthases^6^.

Because F_1_F_o_ ATP synthase can operate in reverse, cells have evolved mechanisms to avoid wasteful hydrolysis of ATP that could occur under physiological conditions. Bacterial ATP synthases appear to utilize a range of different mechanisms for inhibition, with nucleotides, ions and conformational changes likely making contributions^7,8^, with functional studies showing these mechanisms to be mutually exclusive^9^. MgADP is known to inhibit ATPases^10,11^, and can cause the enzyme to fall into a low energy minimum in which MgADP is bound tightly to the catalytic sites. In *E. coli* and other related bacteria, the C-terminal domain of subunit ε (εCTD) also appears to play a crucial role in inhibiting the enzyme by inserting into the F_1_ motor and blocking rotation of the central stalk^7,12-14^. In *E. coli*, the εCTD is comprised of two short helices, residues 86-101 and 110-124 (referred to as εCTH1 and εCTH2 respectively) connected by a linker. Multiple structural studies examining *E. coli* F_1_ or F_1_F_o_ ATP synthase either in the absence of nucleotide^15^ or in the presence of AMPPNP^13^ or MgADP^16^, have shown the εCTD oriented in an extended “up” position, and the isolated subunit has also been crystallized in the condensed “down” position^17^. Our previous ∼5 Å resolution cryo-Electron Microscopy (cryo-EM) study of *E. coli* F_1_F_o_ ATP synthase following incubation with 10 mM MgATP^18^ showed that, under these conditions, the εCTD transitions to a condensed “down” conformation via a “half-up” intermediate in which the εCTH1 remains attached to the central stalk. However, because of the limited resolution of the earlier study, it was not possible to establish the molecular details of how this transition took place.

To understand the detailed structural changes that occur as a result of ATP binding, we have used cryo-EM to examine, at higher resolution, detergent solubilized *E. coli* ATP synthase^19^ following a 45 second incubation with 10 mM MgATP. We show that nucleotide exchange associated with conformational changes of the εCTD and catalytic αβ subunits induces a small rotation of the central stalk in comparison to the structure of the enzyme seen in the presence of MgADP. However, after incubation with MgATP, the β2 site still contains ADP, suggesting that the enzyme is in an ADP inhibited state. Strikingly, the transition of the εCTD to a “down” condensed state in these structures is associated with rotational flexing of the central stalk, which when combined with bending in the peripheral stalk, results in a rotation within the F_o_ motor of two *c* subunits. Truncation constructs of the εCTD show that the potential interaction between the εCTH1 and γ subunit decreases the rate of ATP hydrolysis and aerobic growth. Single molecule^20,21^ and molecular dynamic studies^22^ have indicated that the central rotor could be flexible, but the work presented here shows how this torsional flexibility in the rotor is achieved and how subunit ε is able to modulate it. Furthermore, because the εCTD has been shown to be important for pathogenic bacterial virulence and survival^23-25^ this information may aid the development of bacterial antibiotics targeting these inhibitory mechanisms.

## Results

### Nucleotide occupancy and conformation of the F_1_-ATPase following incubation with MgATP

300 kV cryo-EM was employed to obtain sufficiently high resolution to define how MgATP induces changes within the F_1_ motor. Maps of *E. coli* F_1_F_o_ ATP synthase in the presence of 10 mM MgATP were obtained using methods similar to those in previous studies^15,16,18,26^ (Extended Data Fig. 1 and 2) and provided superior structural information than was observed previously using 200 kV for this complex in the presence of MgATP^18^. The overall resolution improved from 5-6 Å to ∼3 Å, which enabled bound nucleotides to be identified and modeled (Fig. 1a). Previous work^15^ had identified the three major conformational states of the enzyme in which the central stalk is rotated by ∼120° relative to peripheral stalk (termed “State 1”, “State 2” and “State 3” and which refer to the enzyme operating in ATP hydrolysis direction) and the data presented here enabled the generation of cryo-EM maps of these states that had resolutions of 3.0, 2.7 and 3.0 Å, respectively (Extended Data Fig. 2). These maps showed that the nucleotide occupancy and conformation of the F_1_-ATPase differs when the enzyme is incubated with MgATP rather than MgADP^16^. After incubation with 10 mM MgATP, all the non-catalytic α subunits contained MgATP (Extended Data Fig. 3), whereas the contents of the β subunits varied: β1 (β_DP_) contained MgADP, β2 (β_E_) contained ADP and β3 (β_TP_) contained MgATP (Fig. 1a). Compared to the same enzyme imaged in the presence of 10 mM MgADP^16^, the central stalk was rotated ∼10° in the synthase direction (Fig. 1b), the εCTH2 had dissociated from the central stalk, and the β1 (β_DP_) subunit had closed to contact the γ subunit (Fig. 1c). Even with the high-resolutions estimated for these maps, the F_o_ region and position of the ε subunit remained ambiguous (Extended Data Figure 4), hence further data processing was performed to verify the location of the εCTD and *c*-ring, as described in the following section.

**Figure 1:**
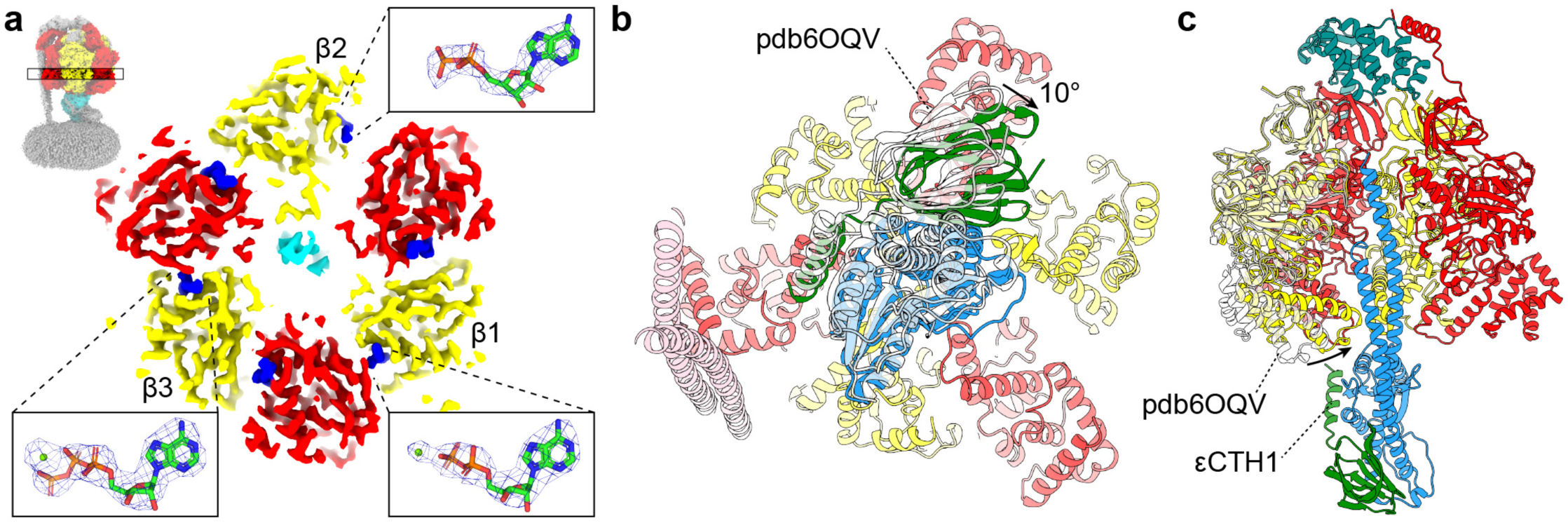
Nucleotide occupancy and conformational changes in F_1_-ATPase following incubation with MgATP. (**a**) Horizontal section of the State 2 *E. coli* F_1_F_o_ ATP synthase cryo-EM map viewed from above, together with details of the catalytic sites in the β subunits (with equivalent mitochondrial F_1_ nomenclature^4^: β1 = β_DP_, β2 = β_E_, β3 = β_TP_^13^). Subunits α in red, β in yellow and γ in cyan, with nucleotide density in dark blue. Higher magnification details of the catalytic sites (inserts) show atomic models, together with cryo-EM maps for the nucleotides (blue mesh). β_DP_ (β1) contains MgADP, β_E_ (β2) contains ADP, and β_TP_ (β3) contains MgATP. Section of map contoured to 0.028 in ChimeraX^27^ and mesh for nucleotides contoured to isolevel 10 in PyMol (Schrödinger). (**b** and **c**) Comparison of the F_1_-ATPase from State 2 *E. coli* F_1_F_o_ ATP synthase after incubation with MgATP (this study; α in red, β in yellow, γ in blue, ε in green, *δ* in teal and b in pink) or MgADP (pdb6OQV^16^; transparent white). (**b**) F_1_-ATPase viewed from below shows that the central stalk (subunits γ and ε) has rotated ∼10° in the clockwise direction after incubation with MgATP. (**c**) F_1_-ATPase viewed from the side, with the closest αβ dimer removed for clarity, shows that the β1 (β_DP_) subunit has closed to contact the γ subunit and the εCTH2 has dissociated from the central stalk after incubation with MgATP (unmodelled as it is not visible in the map).

**Figure 2:**
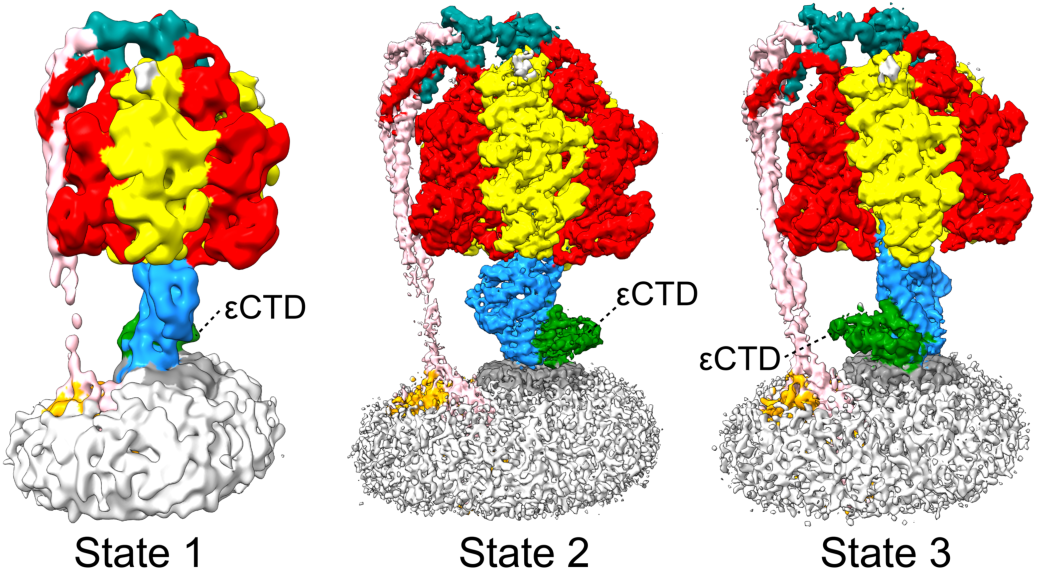
Cryo-EM structures of *E. coli* F_1_F_o_ ATP synthase in three rotational states without εCTD inhibition. Maps of the three rotational states observed by classification on the central stalk (State 1 “down”, State 2 “down” and State 3 “down” in Extended data Fig. 2). Subunits α in red, β in yellow, γ in blue, ε in green, *δ* in teal, *a* in orange, *b* in pink and *c* in grey, with detergent micelle in white. The εCTD (labelled) is in the condensed “down” state in all three maps.

**Figure 3:**
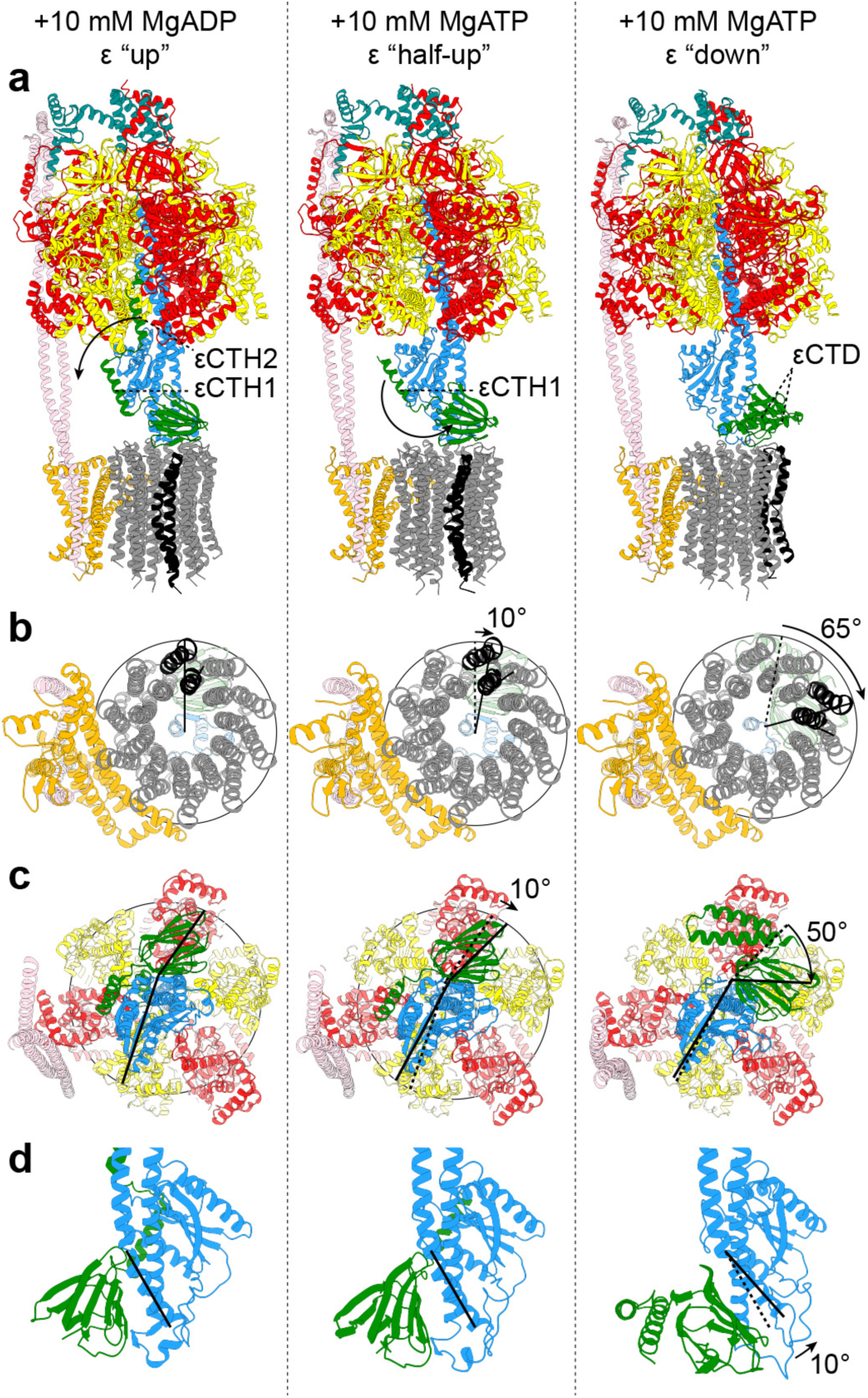
Structural rearrangements of *E. coli* F_1_F_o_ ATP synthase following incubation with ATP. Left; after incubation with 10 mM MgADP (pdb6OQV^16^). Middle; after incubation with 10 mM MgATP where the εCTD is “half-up”. Right; after incubation with 10 mM MgATP where εCTD is “down” (all “State 2” rotary conformation). Subunits α in red, β in yellow, γ in blue, ε in green, *δ* in teal, *a* in orange, *b* in pink and *c* in grey, with a single *c* subunit colored black to highlight rotation of the *c*-ring. **a** and **b** are superposed using the stator *a* subunit, and **c** and **d** are superposed onto the F_1_-ATPase β barrel (residues α26-101 and β1-70). (**a**) Viewed from the side: the εCTD transitions from the “up” conformation, in which the εCTH2 is inserted into the F_1_ motor, to a “down” conformation via a “half-up” intermediate where only the εCTH1 interacts with the central rotor. (**b**) Viewed from below: the *c*-ring rotates two *c* subunit positions (∼75°) between the MgADP structure and MgATP εCTD “down” structure, ∼10° is facilitated by the rotation induced by ATP binding to the catalytic domain and ∼65° is facilitated by the transition of the εCTD from “up” to “down”. (**c**) Viewed from *c*-ring: binding of ATP induces a ∼10° rotation of the central stalk and the εCTD “up” to “down” transition induces a ∼50° rotation of the εNTD which is attached to the γ subunit and *c*-ring (see Fig. 4 also). (**d**) Viewed from the side, zoomed on the central rotor: the “foot” of the γ subunit bends to facilitate the rotation observed between the εCTD “half-up” and “down” states.

**Figure 4:**
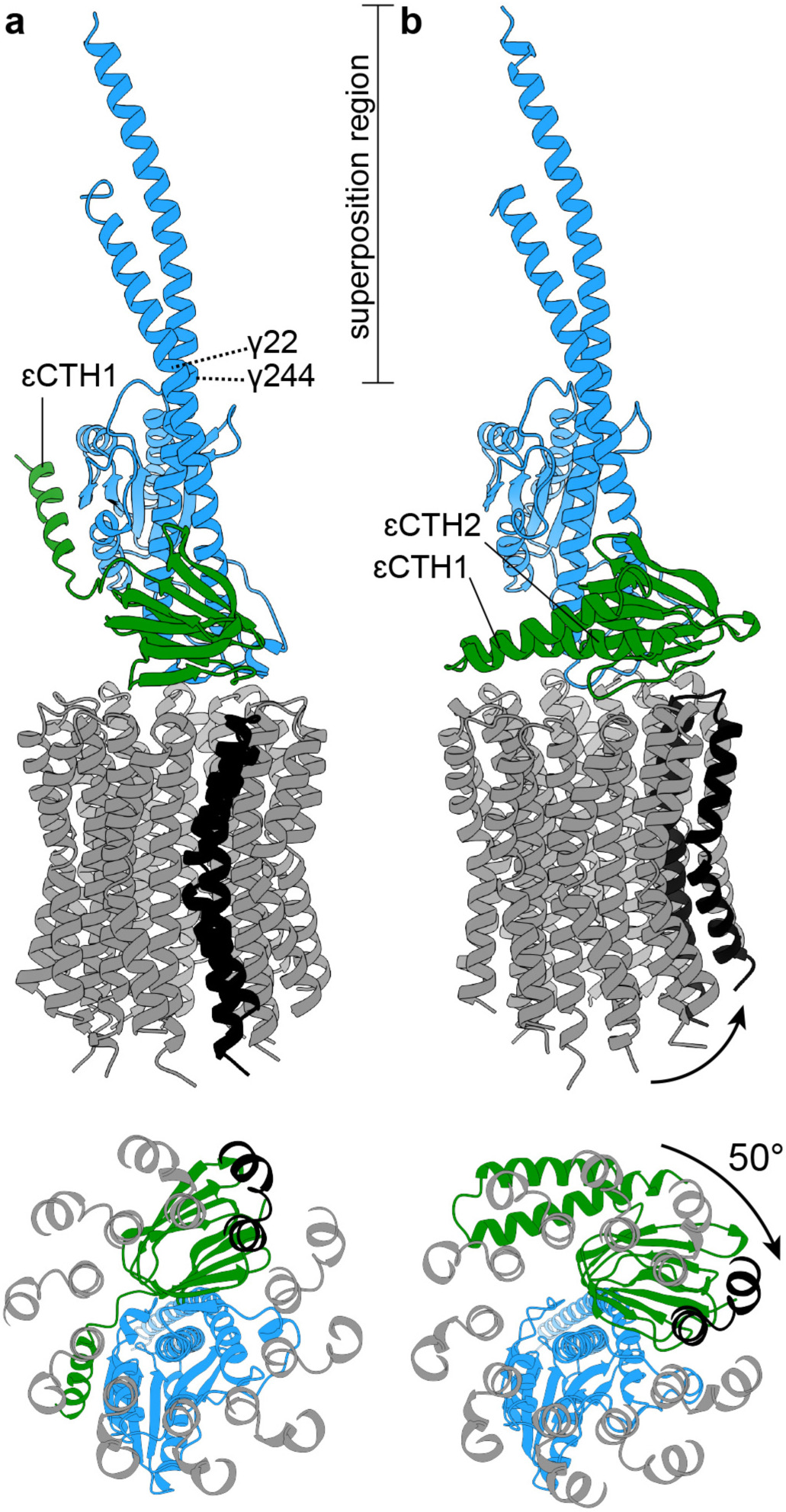
Flexing within central stalk facilitates c-ring to rotation. Superposition on the N- and C-termini of the γ subunit (residues 1-22 and 245-284) of the State 2 “half-up” and “down” central stalk structures. Colors as in Fig. 3. Top; viewed from the side, and bottom; viewed from the periplasm. (**a**) The central rotor of the State 2 “half-up” structure. (**b**) The central rotor of the State 2 “down” structure. The *c*-ring is rotated ∼50° clockwise (synthesis direction) relative to the “half- up” structure.

### Identification of sub-states

Further analysis of the data was implemented using Relion^28^ and identified a series of sub-states that corresponded to different rotational and inhibition states of *E. coli* F_1_F_o_ ATP synthase. Masked 3D classification without image shifts focused on the central rotor highlighted sub-classes in which the εCTD adopted either a condensed “down” conformation or extended “half-up” conformation (Extended Data Fig. 2). The maps generated for each of these sub-states contained detailed information on the position of the εCTD, even though the overall resolution of was reduced (in the range of 3.1-7.2 Å), likely due to the smaller number of particles on which they were based. Three of the maps from this classification describe three rotational positions of *E. coli* F_1_F_o_ ATP synthase with the εCTD in the condensed “down” position, and consequently are not inhibited by this protein motif, thereby allowing rotation of the central stalk (Fig. 2 and Movie 1). However, due to the likely high flexibility of this sample, it was still difficult to unequivocally assign the position of the membrane domain subunits in the maps. To obtain clearer information in this region, a refinement that focused on only the F_o_ motor was performed (Extended Data Fig. 2) and produced maps of sufficient detail to enable fitting of the membrane region (consisting of the *a, b* and *c* subunits) using the structure identified from the same sample imaged with MgADP^16^.

### The εCTD attenuates the central stalk

In the previous lower-resolution cryo-EM study, it was only possible to observe the different conformations of the εCTD by comparing structures from different rotational states (State 1 showed εCTD in the “half-up” conformation, State 2 showed a blurred εCTD and State 3 showed the εCTD in the “down” state). In this study, we were able to observe the εCTD in two conformations in both State 1 and State 2, whereas State 3 was only observed in the “down” state (Extended Data Fig. 2). The increased number of states identified was likely a result of the higher resolution obtained by using a 300 kV accelerator voltage to collect the images together with the larger number of particles in the dataset. The importance of the number of particles was also underlined by the reduced resolution obtained as a consequence of having smaller numbers of particles corresponding to εCTD “down” sub-states in State 1 (Extended Data Fig. 2). By contrast, State 2 contained a more similar number of particles for both the εCTD “half-up” and “down” states, and consequently the maps were more detailed. When the εCTD “half-up” and “down” structures from State 2 were superposed using the *a* subunit of the stator, clear differences between the rotational positions of the rotor *c*-ring were observed (Fig. 3a and b). Most interestingly, the *c*- ring was rotated ∼65° in the synthesis direction as the εCTD transitioned from the “half-up” to the “down” sub-state, with this rotation facilitated by flexing in both the peripheral and central stalks.

The different states showed evidence for torsional flexing in the central stalk that originated from a rotation of the ε subunit relative to the γ subunit (Fig. 3c and 4), with the flexing facilitated near the “foot” of subunit γ (Fig. 3d). In the isolated rotor, the rotation observed between the εCTD “half-up” and “down” states was ∼50° (Figure 4). This movement is prevented in the “half-up” conformation by the εCTH1 binding to the opposite side of the γ subunit to which the ε N-terminal domain (εNTD) binds, suggesting that the subunit ε may regulate the flexibility of the γ subunit and thereby the efficiency of the enzyme. To test this hypothesis, two ε subunit truncation mutants were generated: the first removed εCTH2 (εΔCTH2; residues ε1-104) and second removed both εCTH1 and εCTH2 (εΔCTH1+2; residues ε1-81) (Fig. 5a). ATP regeneration assays showed that, although the εΔCTH2 mutant had higher turnover than wildtype enzyme, the εΔCTH1+2 mutant showed even higher turnover (Fig. 5b), indicating that the enzyme had higher activity when the central stalk is free from restriction by the εCTD. Aerobic growth assays of these same mutants showed that removal of the εCTH2 induced a growth phenotype, (Fig. 5c) similar to that seen when the five C-terminal residues are deleted from the ε subunit^29^.

**Figure 5:**
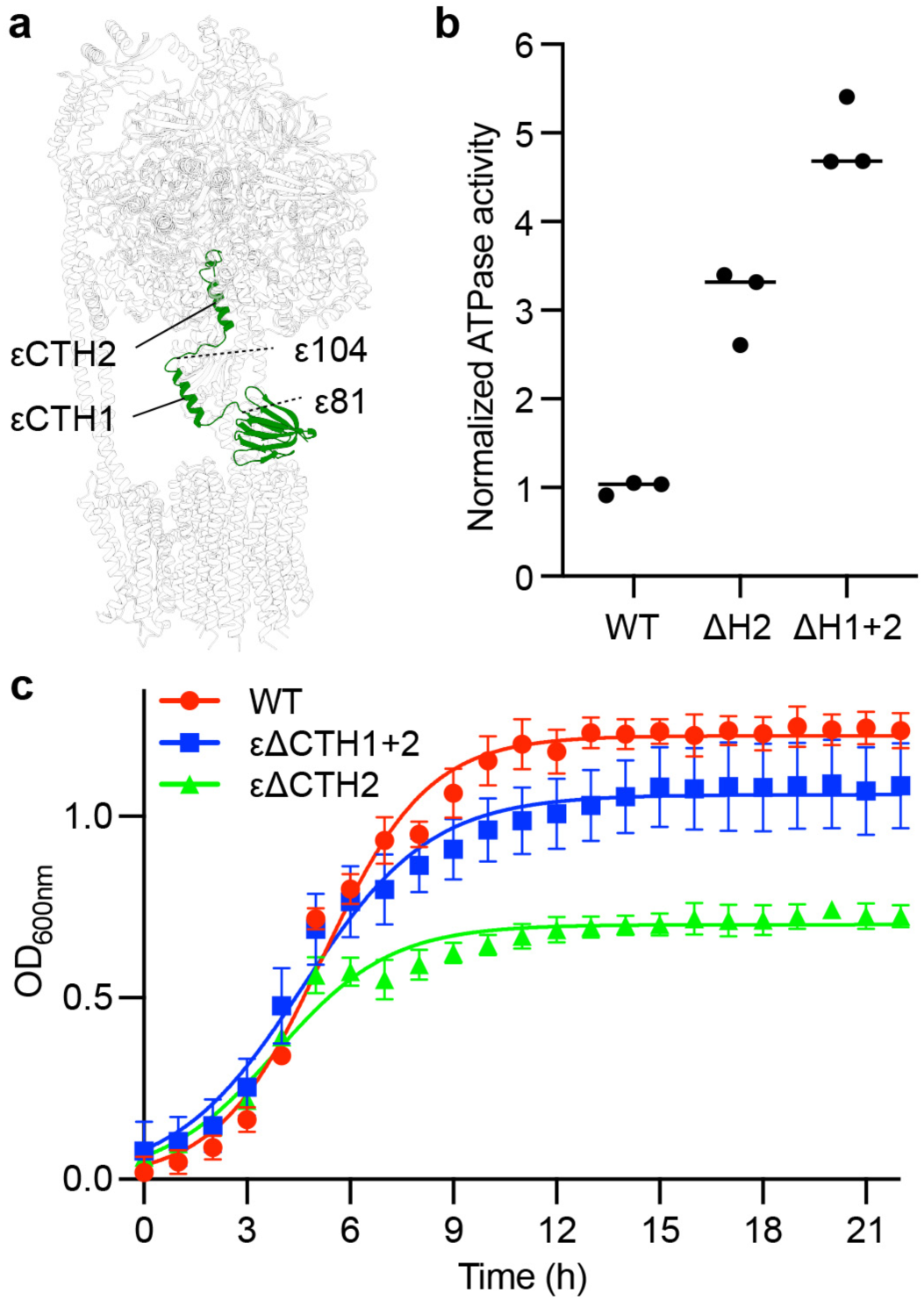
The εCTD attenuates ATPase activity and aerobic growth beyond binding with the catalytic subunits. (**a**) Location of residues ε81 and ε104 in the intact F_1_F_o_ enzyme. (**b**) ATP regeneration assays of WT (containing full length subunit ε), εΔCTH2 (ε1-104; with the εCTH2 and linker removed) and εΔCTH1+2 (ε1-81; with εCTH1, εCTH2 and linkers removed). All data points and mean shown (normalized to the mean of WT). Removal of εCTH2 results in higher ATP turnover than wild-type. Removal of both εCTH1 and εCTH2 shows higher ATP turnover than just removing εCTH2. (**c**) Succinate growth assays show that removal of εCTH2 reduces *E. coli* aerobic growth. n of 3; mean and standard deviation shown for each time point; measurements taken on distinct samples; line calculated using logistic growth analysis in Prism 8 for macOS. εΔCTH1+2 grows to a lower density, suggesting that the flexibility of the central stalk plays a role in cellular ATP synthase function.

## Discussion

The cryo-EM and functional studies presented here provide new information on how conformational changes are induced in *E. coli* ATP synthase by MgATP and indicate how subunit ε is able to modulate the function of the enzyme.

The cryo-EM structures provide additional detailed information on the structural changes and nucleotide occupancy introduced by binding MgATP and indicate that the transition of the εCTD from an “up” to a “down” state via a “half-up” intermediate is coupled with movements within both the F_1_ and F_o_ motors. Compared to the same enzyme observed after incubation with MgADP, incubation with MgATP induces the β1 (β_DP_) subunit in F_1_ to close and contact the γ subunit with P_i_ being liberated from the active site. β3 (β_TP_) exchanges ADP for ATP and the γ subunit rotates ∼10° in the synthesis direction. The β2 (β_E_) site is still loosely occupied by the product (ADP), similar to that seen in the bovine enzyme crystalized with reduced Mg^2+^ concentration (termed F1–PH)^30^ (Fig. 1a). Given that we observe MgADP without P_i_ in β1 (β_DP_), we hypothesize that the structure we observed in this study represents the enzyme paused in the MgADP inhibited state, where P_i_ has been released but MgADP is bound to the β1 (β_DP_) site. ATPase assays previously performed on this exact protein purification incubated with 10 mM MgATP for 45 seconds, predicted that the enzyme is being imaged in the presence of ∼9.75 mM ATP and ∼0.3 mM ADP^18^. So although the enzyme is undergoing ATP hydrolysis, it is likely paused in the MgADP inhibited state, consistent with the pausing observed in single molecule studies on related F_1_-ATPase enzymes^31^.

Although flexible coupling between the F_1_ and F_o_ motors is necessary to facilitate efficient enzyme function, whether this flexibility originates from the peripheral or central stalk has been controversial^20,22,32-35^. To date structural studies have only shown flexibility within the peripheral stalk^36^, with a central stalk remaining essentially rigid in all rotational substes^14,16,37-40^. Our previous work on the *E. coli* enzyme incubated with MgADP showed that the peripheral stalk is able to flexibly couple the F_1_ and F_o_ motors, facilitating a single sub-step in the F_o_ motor without any flexing in the central stalk. The structural and function data obtained here on the *E. coli* enzyme shows that flexibility also stems from the central stalk. Single molecule^20,21^ and molecular dynamic studies^22^ have indicated that the rotor can be flexible, but the present work shows how this flexibility is conferred and indicates that rotor torsional flexibility is important *E. coli* function. We hypothesize that the bridging of the εNTD to the opposite side of subunit γ, as seen in the “half-up” state, enables the εCTD is able to attenuate the rotational flexibility of the central stalk and thereby change the efficiency of the enzyme in response to cellular conditions. Because, in other species, the εCTD has been shown to bind ATP selectively^41-43^, cellular ATP concentration is likely to be the signal that controls this function of the subunit. The maps presented in this study are consistent with ATP binding to the εCTD when it is in the “down” position (Extended Data Fig. 5). However, the maps are not sufficiently detailed to distinguish between ATP and ADP, or even other small molecules. In the related bacteria *Geobacillus stearothermophilus* (more commonly termed *Bacillus* PS3), the εCTD in the “up” conformation forms a single continuous helix^14,44^ that does not bridge the γ subunit and εNTD in the same manner as *E. coli* enzyme. Therefore, it is unlikely that bacteria such as *Bacillus* PS3 utilize this mechanism to attenuate the flexibility of the central stalk.

The transition of the εCTD to the “down” state shown in this study corresponds to the *c*- ring being rotated ∼65° in the synthase direction relative to the stator. However, this rotation is a combination of movements in the peripheral and central stalks, with ∼15° facilitated by flexing of the peripheral stalk and ∼50° facilitated by twisting of the central stalk. The rotation of the *c*-ring between the εCTD “up” and “down” states is in the synthesis direction (clockwise when viewed from the periplasm), but because the enzyme was frozen in the presence of ATP (and therefore rotating in the hydrolysis direction), this rotation corresponds to the *c*-ring being retarded in these states. This observation highlights the resistance likely incurred at the stator rotor interface or between the *c*-ring and lipids. The molecular events that occur between the autoinhibited and MgADP states can be appreciated by interpolating between rotational State 2 of the enzyme incubated with MgADP^16^ and the half-up and down substates of rotational State 2 incubated with MgATP (Fig. 3 and Movie 2). These structures indicate that the εCTH2 first dissociates from the catalytic head and the β1 (β_DP_) subunit releases P_i_. The β1 (β_DP_) subunit then closes to the MgADP inhibited state, rotating the γ subunit ∼10°. In this state, the εCTH1 is still bound to the γ subunit, but this can dissociate and bind nucleotide with the rest of the ε subunit to generate the down state. The reduced contacts between the γ and ε subunits generated in this way increase the flexibility of the central stalk, which can facilitate flexible coupling between the F_1_ and F_o_ motors, smoothing the process in an analogous way to the fluid coupling method found in automatic gear boxes.

## Data availability

The models generated and analyzed during the current study are available from the RCSB PDB: 7KA5, 7KA6, 7KA7, 7KA8, 7KA9

The cryo-EM maps used to generate models are available from the EMDB: 22711, 22759, 22760, 22761, 22762, 22763, 22764, 22765, 22766, 22767, 22768, 22769

## Acknowledgments

We wish to thank and acknowledge Dr Craig Yoshioka and Dr Claudia López (Oregon Health & Sciences University (OHSU)) for data collection and processing expertise. A.G.S was supported by a National Health and Medical Research Council Fellowship APP1159347 and Grant APP1146403. We wish to thank and acknowledge the use of the Victor Chang Innovation Centre, funded by the NSW Government, and the Electron Microscope Unit at UNSW Sydney, funded in part by the NSW Government. A portion of this research was supported by NIH grant U24GM129547 and performed at the Pacific Northwest Centre for Cryo-EM at OHSU and accessed through EMSL (grid.436923.9), a DOE Office of Science User Facility sponsored by the Office of Biological and Environmental Research. Molecular graphics and analyses performed with UCSF ChimeraX, developed by the Resource for Biocomputing, Visualization, and Informatics at the University of California, San Francisco, with support from National Institutes of Health R01-GM129325 and the Office of Cyber Infrastructure and Computational Biology, National Institute of Allergy and Infectious Diseases

## Contributions

M.S., R.I and A.G.S. conceived the study and wrote the manuscript. M.S. performed the formal analysis of the study. JLW and YCZ aided in data acquisition, analysis and interpretation. A.G.S. supervised the study.

## Competing interests

Authors declare no competing interests

## Methods

Protein purification: The *E. coli* F_1_F_o_ ATP synthase protein was prepared as described in Sobti et al. 2019^18,26^. Cysteine-free *E. coli* ATP synthase (all cysteines residues substituted with alanine and a His-tag introduced on the β subunit) was expressed in *E. coli* DK8 strain^19^. Cells were grown at 37°C in LB medium supplemented with 100 μg/ml ampicillin for 5 h. The cells were harvested by centrifugation at 5,000 g, providing ∼1.25 g cells per litre of culture. Cells were resuspended in lysis buffer containing 50 mM Tris/Cl pH 8.0, 100 mM NaCl, 5 mM MgCl_2_, 0.1 mM EDTA, 2.5% glycerol and 1 μg/ml DNase I, and processed with three freeze thaw cycles followed by one pass through a continuous flow cell disruptor at 20 kPSI. Cellular debris was removed by centrifuging at 7,700 × g for 15 mins, and the membranes were collected by ultracentrifugation at 100,000 × g for 1 h. The ATP synthase complex was extracted from membranes at 4°C for 1 h by resuspending the pellet in extraction buffer consisting of 20 mM Tris/Cl, pH 8.0, 300 mM NaCl, 2 mM MgCl_2_, 100 mM sucrose, 20 mM imidazole, 10% glycerol, 4 mM digitonin and EDTA-free protease inhibitor tablets (Roche). Insoluble material was removed by ultracentrifugation at 100,000 g for 30 min. The complex was then purified by binding on Talon resin (Clontech) and eluted in 150 mM imidazole, and further purified with size exclusion chromatography on a 16/60 Superose 6 column equilibrated in a buffer containing 20 mM Tris/Cl pH 8.0, 100 mM NaCl, 1 mM digitonin and 2 mM MgCl_2_. The purified protein was then concentrated to 11 μM (6 mg/ml), and snap frozen and stored for grid preparation.

Mutant constructs were made using the following primers:

εΔCTH2: Forward primer 5’-aagcgaaacgtaaggctgaagagcactaacaccggcttgaaaagcacaaa-3’ Reverse primer 5’-tggcttttgtgcttttcaagccggtgttagtgctcttcagccttacgttt-3’

εΔCTH1+2: Forward primer 5’-aacgtgaccgttctggccgactaacaccggcttgaaaagcacaaa-3’ Reverse primer 5’-ggcttttgtgcttttcaagccggtgttagtcggccagaacggtcacgtt-3’

Cryo-EM grid preparation: 1 μl of 100 mM ATP/100 mM MgCl_2_ (pH 8.0) was added to an aliquot of 9 μl of purified cysteine-free *E. coli* F_1_F_o_ ATP synthase at 11 μM (6 mg/ml) and the sample was incubated at 22°C for 30 s, before 3.5 μl was placed on glow-discharged holey gold grid (UltrAufoils R1.2/1.3, 200 Mesh). Grids were blotted for 3 s at 22°C, 100 % humidity and flash-frozen in liquid ethane using a FEI Vitrobot Mark IV (total time for sample application, blotting and freezing was 45 s).

Data collection: Grids were transferred to a Thermo Fisher Talos Arctica transmission electron microscope (TEM) operating at 200 kV and screened for ice thickness and particle density. Suitable grids were subsequently transferred to a Thermo Fisher Titan Krios TEM operating at 300 kV equipped with a Gatan BioQuantum energy filter and K3 Camera at the Pacific Northwest Centre for Cryo-EM at OHSU. Images were recorded automatically using serial EM at 81,000 x magnification yielding a pixel size of 0.54 Å (K3 operating in super resolution mode). A total dose of 48 electrons per Å^2^ was used spread over 77 frames, with a total exposure time of 3.5 s. 8,620 movie micrographs were collected (Extended Data Fig.1).

Data processing: MotionCorr2^45^ was used to correct local beam-induced motion and to align resulting frames, with 9×9 patches and binning by a factor of two. Defocus and astigmatism values were estimated using Gctf^38^ and 8,215 micrographs were selected after exclusion based on ice contamination, drift and astigmatism. ∼1,000 particles were manually picked and subjected to 2D classification to generate templates for template picking in cryoSPARC^46^, yielding 869,147 particles. These particles were binned by a factor of four and subjected to 2D classification generating a final dataset of 429,638 particles. The locations of these particles were then imported into Relion^28^, re-extracted at full resolution, and further classified into 3D classes using a low pass filtered cryo-EM model generated from a previous study^15^, yielding the three main states related by a rotation of the central stalk (State1, State2 and State3 with 100,831, 215,003, and 113,804 particles, respectively). Focused classification, using a mask comprising the lower half of the central rotor, was implemented without performing image alignment in Relion 3.0, yielding the “half-up” and the “down” sub-classes in each of the three main states. A further F_o_ focused classification without image shifts was performed on each of the “half-up” and “down” sub-classes to elucidate the position of F_o_ subunits in the respective sub-states. See Extended Data Fig. 2 for a flowchart describing this classification, Extended Data Table 1 for a summary of data collection/processing statistics and Extended Data Fig. 6 for FSC curves.

Model building: Models were built and refined in Coot^47^, PHENIX^48^ and ISOLDE^49^ using pdbs 6OQT, 6OQV, 6OQW^16^ (*E. coli* ATP synthase incubated with MgADP) and 1AQT^17^ (isolated *E. coli* ATP synthase subunit ε) as guides. See Extended Data Table 1 for a summary of refinement and validation statistics.

ATP regeneration assays: ATP regeneration assays were performed as in Sobti et al 2020^26^. In short; 10 μg of protein was added to 100 mM KCl, 50 mM MOPS pH 7.4, 1 mM MgCl2, 1 mM ATP, 2 mM PEP, 2.5 units/ml pyruvate kinase, 2.5 units/ml lactate dehydrogenase and 0.2 mM NADH, and monitored for OD at 340 nm at for up to 20 min (Extended Data Fig. 7).

Phenotypic assay for respiratory growth: Aerobic growth assays were performed similarly Shah & Duncan 2015^29^. Briefly, single colonies of the three constructs (WT, εΔCTH2 and εΔCTH1+2) transformed into DK8 cells^19^ were grown at 37°C in 10ml of LB + 100 μg /ml ampicillin until OD at A_600_ was ∼0.4. The cells were then diluted 40-fold into minimal medium supplemented with 1mM MgCl_2_, 0.1% amino acid solution, 0.1% trace elements, 100 μg /ml ampicillin and 0.8% succinate. 0.4 ml cultures were set up in triplicates in a 48-well transparent plate sealed with a Breathe-Easy® sealing membrane. Growth was monitored and measured at 37°C in a PHERAstar FS plate reader with shaking at 100 rpm until cells reached stationary phase.

## Notes

### Competing Interest Statement

The authors have declared no competing interest.

