## SUPPLEMENTARY INFORMATION for "Subunit epsilon of *E. coli* F_1_F_o_ ATP synthase attenuates enzyme activity by modulating central stalk flexibility"

Meghna Sobti et al.

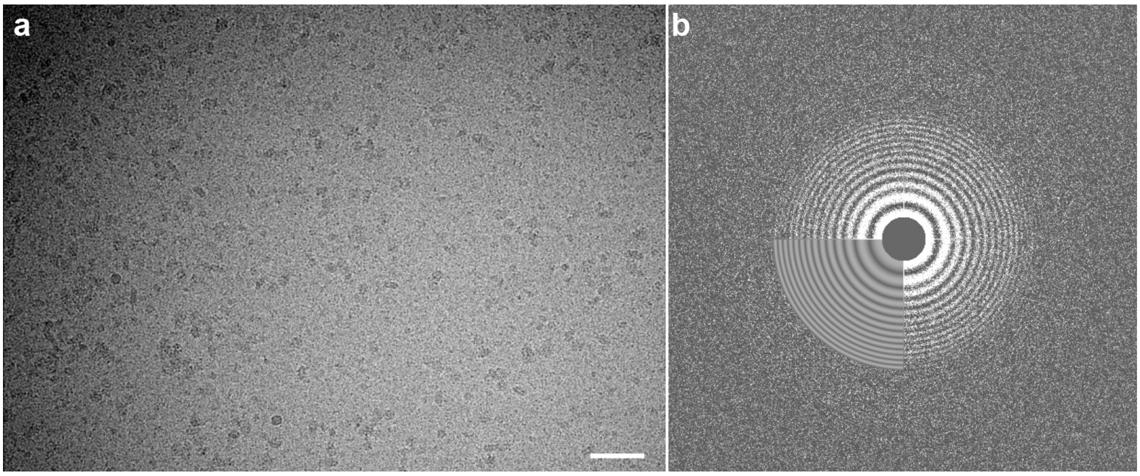


**Extended Data Fig. 1:** Cryo-EM micrograph and power spectrum. (**a**) Representative micrograph showing ATP synthase particles (white scale bar is equivalent to 50 nm) and (b) Gctf[1] (implemented in cryosparc[2]) output of same micrograph. The data was taken on a distinct sample, we have imaged *E. coli* ATP synthase many times and taken full cryo-EM datasets >10 times. The micrograph shown was chosen to show clear ATP synthase particles (high defocus) and is devoid of ice contamination and particle aggregation.

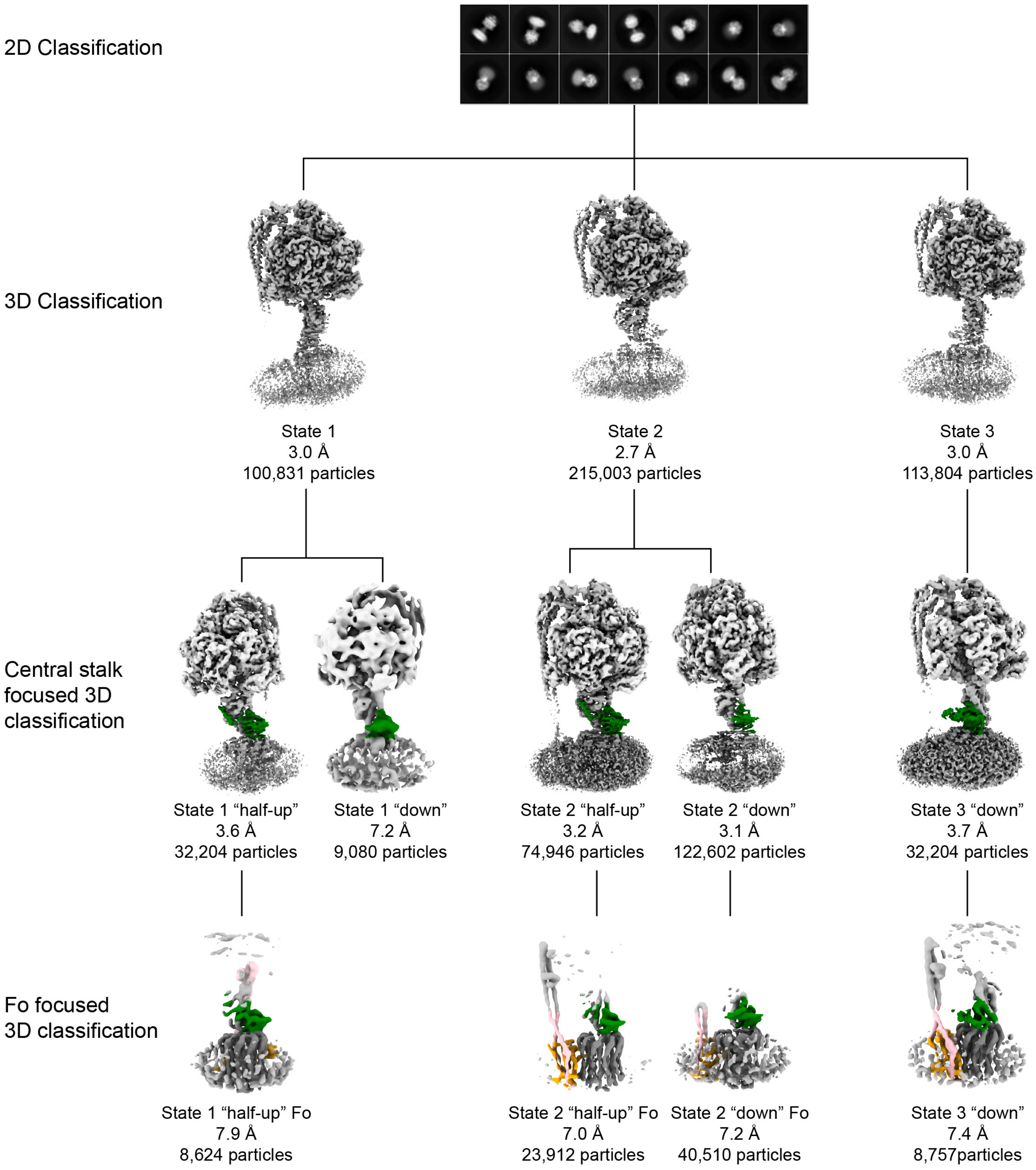


**Extended Data Fig. 2: Flowchart describing the classification of particles into distinct conformations.**

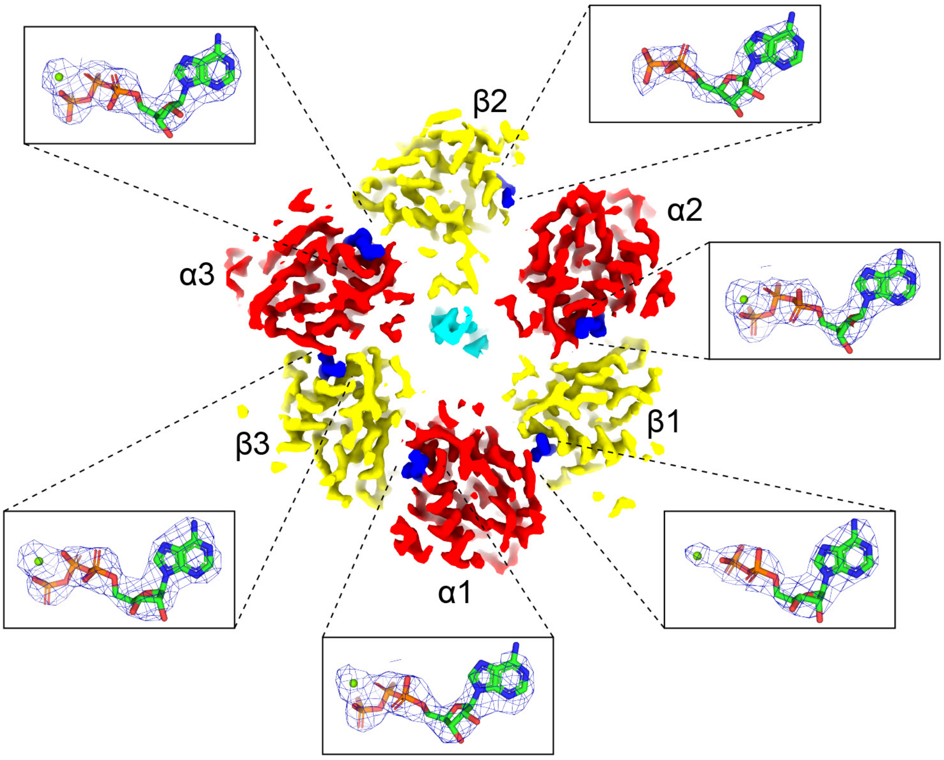


**Extended Data Fig. 3: $F_1$-ATPase nucleotide assignment.** Horizontal section of the State 2 *E. coli* $F_1F_o$ ATP synthase cryo-EM map (2.7 Å resolution) viewed from above (as in Fig. 2), together with details of the catalytic site in the α and β subunits (with equivalent Mitochondrial $F_1$ nomenclature[3]: 1 = DP, 2 = E, 3 = TP[4]). Subunits α in red, β in yellow and γ in cyan, with nucleotide density in blue. Higher magnification details of the catalytic sites (inserts) show atomic models, together with cryo-EM maps for the nucleotides (blue mesh). All α subunits contain MgATP. $\beta_{DP}$ (β1) contains MgADP, and $\beta_E$ (β2) contains ADP and $\beta_{TP}$ (β3) contains MgATP. Section of map contoured to 0.028 in ChimeraX[5] and mesh for nucleotides contoured to isolevel 10 in PyMol (Schrödinger)

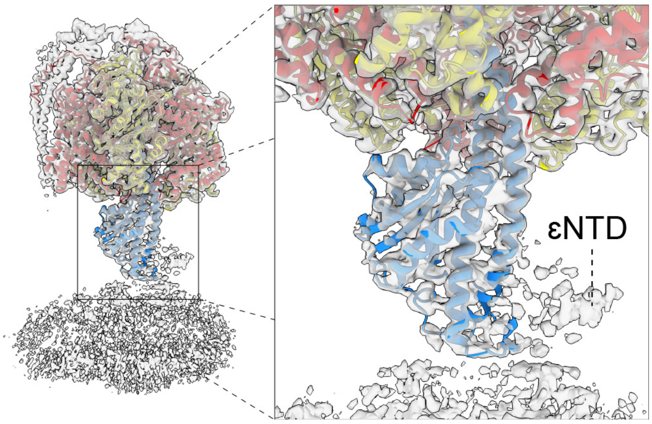


**Extended Data Fig. 4: The positions of the $F_o$ motor and ε subunit are not evident in the initial 3D classification.** The cryo-EM maps did not show interpretable density for the $F_o$ or ε subunit after initial 3D classification. (**a**) Complete map with $F_1$ atomic model fitted and (**b**) close up of the expected location of subunit ε. "State 2" shown (see Extended Data Fig. 2).

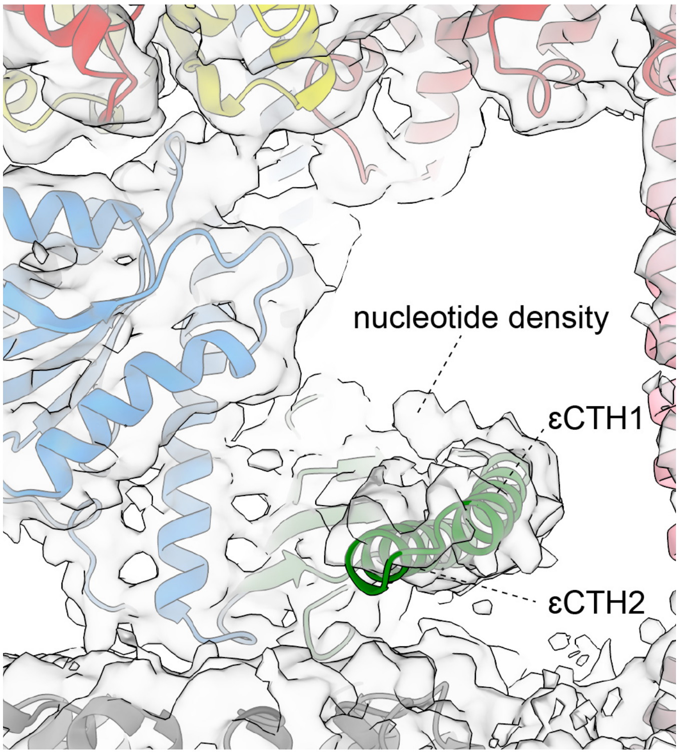


**Extended Data Fig. 5: Density corresponding to a potential small molecule bound to the εCTD.** Cryo-EM map of State 3 "down" shows extra density at the predicted εCTD binding site. The size and shape are consistent with ATP, but the density is not clear enough to unequivocally assign this density.

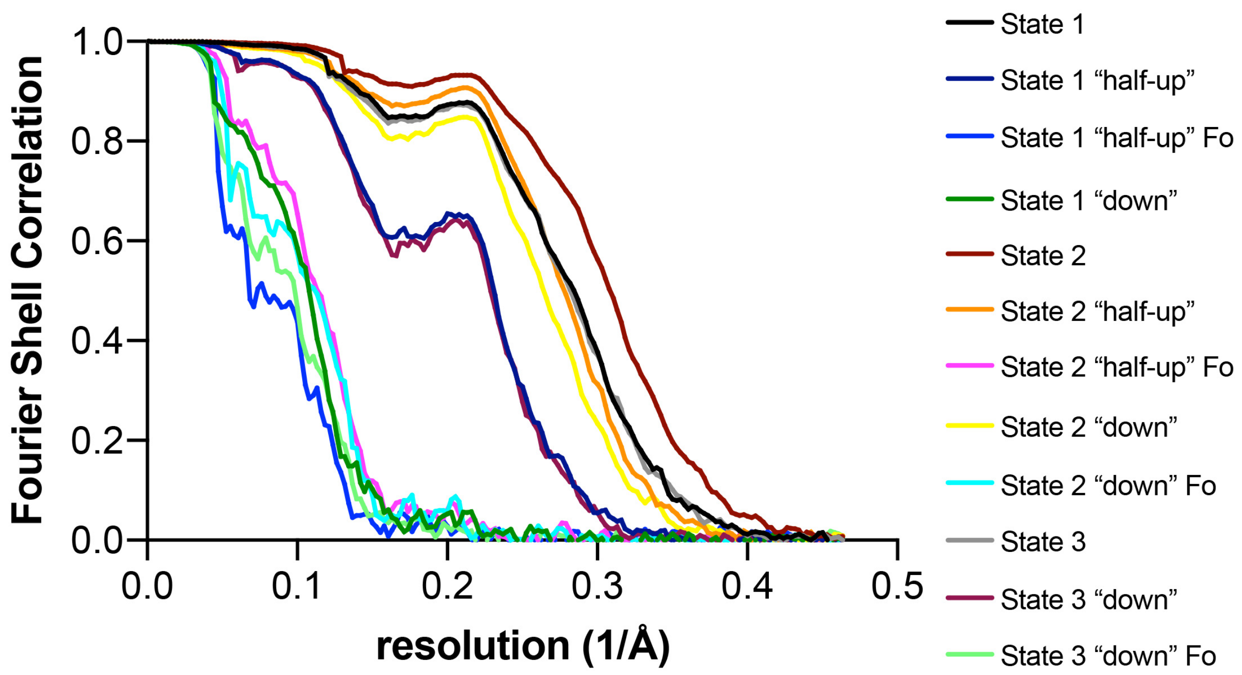


**Extended Data Fig. 6: Fourier shell correlation curves for states shown in Extended Data Fig. 2.** Masked FSC curves from Relion[6] postprocess.

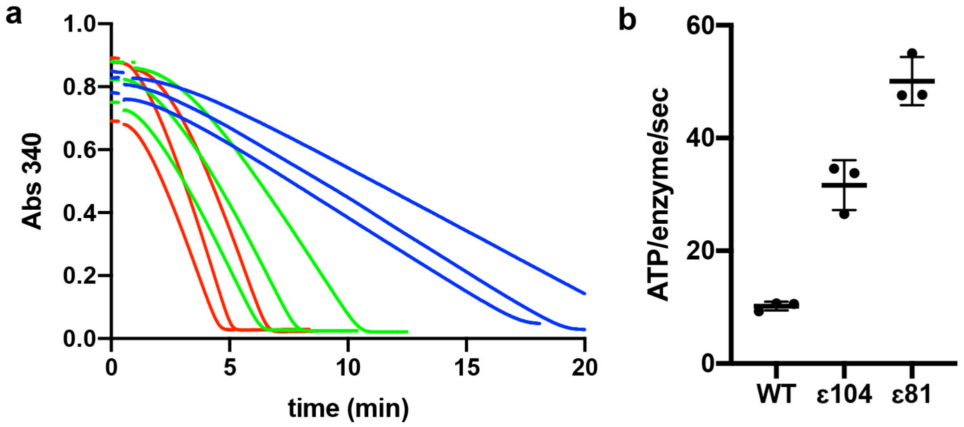


**Extended Data Fig. 7: ATP regeneration assays.** ATP regeneration assays of WT, εΔCTH2 and εΔCTH1+2 mutations. (**a**) Raw traces and (**b**) ATP turnover per enzyme per second (mean and standard deviation shown.

### Extended Data Table 1: Cryo-EM data collection, refinement and validation statistics.

| | #1 State1 (EMD-22711) | #2 State1 half-up (EMD-22759) (PDB 7KA5) | #3 State1 half-up Fo (EMD-22760) | #4 State1 down (EMD-22768) (PDB 7KA8) | #5 State2 (EMD-22761) | #6 State2 half-up (EMD-22769) (PDB 7KA9) | #7 State2 half-up Fo (EMD-22762) | #8 State2 down (EMD-22763) (PDB 7KA6) | #9 State2 down Fo (EMD-22764) | #10 State3 (EMD-22765) | #11 State 3 down (EMD-22766) (PDB 7KA7) | #12 State 3 down Fo (EMD-22767) |
|---|---|---|---|---|---|---|---|---|---|---|---|---|
| **Data collection and processing** | | | | | | | | | | | | |
| Magnification | 81,000 | 81,000 | 81,000 | 81,000 | 81,000 | 81,000 | 81,000 | 81,000 | 81,000 | 81,000 | 81,000 | 81,000 |
| Voltage (kV) | 300 | 300 | 300 | 300 | 300 | 300 | 300 | 300 | 300 | 300 | 300 | 300 |
| Electron exposure ($e^{-}$/Å$^2$) | 48 | 48 | 48 | 48 | 48 | 48 | 48 | 48 | 48 | 48 | 48 | 48 |
| Defocus range (μm) | 0.8-3.5 | 0.8-3.5 | 0.8-3.5 | 0.8-3.5 | 0.8-3.5 | 0.8-3.5 | 0.8-3.5 | 0.8-3.5 | 0.8-3.5 | 0.8-3.5 | 0.8-3.5 | 0.8-3.5 |
| Pixel size (Å) | 1.08 | 1.08 | 1.08 | 1.08 | 1.08 | 1.08 | 1.08 | 1.08 | 1.08 | 1.08 | 1.08 | 1.08 |
| Symmetry imposed | C1 | C1 | C1 | C1 | C1 | C1 | C1 | C1 | C1 | C1 | C1 | C1 |
| Initial particle images (no.) | 429,638 | 429,638 | 429,638 | 429,638 | 429,638 | 429,638 | 429,638 | 429,638 | 429,638 | 429,638 | 429,638 | 429,638 |
| Final particle images (no.) | 100,831 | 32,204 | 8,624 | 9,080 | 215,003 | 74,946 | 23,912 | 122,602 | 40,510 | 113,804 | 32,204 | 8,757 |
| Map resolution (Å) 0.143 FSC threshold | 3.0 | 3.6 | 7.9 | 7.2 | 2.7 | 3.2 | 7.0 | 3.1 | 7.2 | 3.0 | 3.7 | 7.4 |
| **Refinement** | | | | | | | | | | | | |
| Initial models used (PDB codes) | - | 6OQT | - | 6OQT, 1AQT | - | 6OQV | - | 6OQV, 1AQT | - | - | 6OQW, 1AQT | - |
| Model resolution (Å) 0.5 FSC threshold, masked/ unmasked | - | 3.6/4.1 | - | 8.2/9.0 | - | 3.2/3.6 | - | 3.2/3.4 | - | - | 3.7/4.2 | - |
| Map sharpening *B* factor (Å$^2$) | -57 | -62 | -169 | -125 | -53 | -61 | -185 | -63 | -199 | -58 | -74 | -176 |
| Model composition | | | | | | | | | | | | |
| Non-hydrogen atoms | - | 36,632 | - | 36,881 | - | 36,674 | - | 36,933 | - | - | 36,856 | - |
| Protein residues | - | 4,800 | - | 4,833 | - | 4,814 | - | 4,847 | - | - | 4,837 | - |
| Ligands | - | 11 | - | 11 | - | 11 | - | 11 | - | - | 11 | - |
| *B* factors (Å$^2$) | | | | | | | | | | | | |
| Protein | - | 148 | - | 606 | - | 161 | - | 222 | - | - | 146 | - |
| Ligand | - | 88 | - | 611 | - | 86 | - | 86 | - | - | 65 | |
| R.m.s. deviations | | | | | | | | | | | | |
| Bond lengths (Å) | - | 0.01 | - | 0.009 | - | 0.01 | - | 0.01 | - | - | 0.01 | - |
| Bond angles (°) | - | 1.164 | - | 1.339 | - | 1.256 | - | 1.24 | - | - | 1.2 | - |
| Validation | | | | | | | | | | | | |
| MolProbity score | - | 1.12 | - | 1.90 | - | 1.12 | - | 1.09 | - | - | 1.51 | - |
| Clashscore | - | 2.16 | - | 8.32 | - | 2.19 | - | 1.90 | - | - | 3.58 | - |
| Poor rotamers (%) | - | 1.13 | - | 1.91 | - | 0.45 | - | 0.52 | - | - | 1.86 | - |
| Ramachandran plot | | | | | | | | | | | | |
| Favored (%) | - | 97.58 | - | 96.47 | - | 97.36 | - | 97.25 | - | - | 97.1 | - |
| Allowed (%) | - | 2.36 | - | 3.36 | - | 2.58 | - | 2.71 | - | - | 2.86 | - |
| Disallowed (%) | - | 0.06 | - | 0.17 | - | 0.06 | - | 0.04 | - | - | 0.04 | - |
